# Nonsynonymous A-to-I RNA editing contributes to burden of deleterious missense variants in healthy individuals

**DOI:** 10.1101/462390

**Authors:** Te-Lun Mai, Trees-Juen Chuang

## Abstract

Adenosine-to-inosine (A-to-I) RNA editing is a very common post-transcriptional modification that can lead to A-to-G changes at the RNA level and compensate for G-to-A genomic changes to a certain extent. It has been shown that each healthy individual can carry dozens of missense variants predicted to be severely deleterious. Why strongly detrimental variants are preserved in a population and not eliminated by negative natural selection remains mostly unclear. Here we ask if RNA editing correlates with the burden of deleterious A/G polymorphisms in a population. Integrating genome and transcriptome sequencing data from 447 human lymphoblastoid cell lines, we show that nonsynonymous editing activities (prevalence/level) are negatively correlated with the deleteriousness of A-to-G genomic changes and positively correlated with that of G-to-A genomic changes within the population. We find a significantly negative correlation between nonsynonymous editing activities and allele frequency of A within the population. This negative editing-allele frequency correlation is particularly strong when editing sites are located in highly important genes/loci. Examinations of deleterious missense variants from the 1000 genomes project further show a significantly higher mutational burden in G-to-A changes than in other types of changes. The level of the mutational burden in G-to-A changes increases with increasing deleterious effects of the changes. Moreover, the deleteriousness of G-to-A changes is significantly positively correlated with the percentage of binding motif of editing enzymes at the variants. Overall, we show that nonsynonymous editing contributes to the increased burden of G-to-A missense mutations in healthy individuals, expanding RNA editing in pathogenomics studies.

## INTRODUCTION

Mutations are the ultimate driving force of evolution, providing the major source of genetic variability within a population. Generally, most mutations at functionally important loci, especially those that cause nonsynonymous changes (missense substitutions), are destined for selective elimination because of their deleteriousness. However, it was observed that each healthy individual could carry hundreds of missense substitutions, some of which were homozygous and predicted to be severely deleterious or disease-causing (MacArthur et al. 2012; Tennessen et al. 2012; Xue et al. 2012). Recent analysis of the Human Gene Mutation Database (HGMD^®^) (Stenson et al. 2009) also revealed that 5,132 out of 92,331 missense variants were classified as disease-causing mutations (Stenson et al. 2017). For pathogenomics studies, deleterious variants are often observed in well-established disease-associated genes in population controls, making it difficult to extract pathogenic variants (MacArthur et al. 2012). Many studies have investigated the burden of deleterious variants, or the so-called mutation load, carried by a population and indicated that the persistence of deleterious variants in a population is primarily affected by the strength of genetic drift and negative selection (Kimura et al. 1963; King and Jukes 1969; Lynch and Gabriel 1990; Ohta and Gillespie 1996; Lohmueller 2014; Henn et al. 2015). It is understandable that mildly deleterious variants contribute more to mutation load than severely deleterious ones because the former are subject to relatively weaker selective constraints than the latter (Lohmueller 2014; Henn et al. 2015). However, the reason that strongly detrimental variants (particularly those in the homozygous state) are preserved in a population and not eliminated by negative natural selection remains mostly unclear.

Adenosine-to-inosine (A-to-I) RNA editing is a very common mechanism in metazoans (Porath et al. 2017a; Porath et al. 2017b; Hung et al. 2018). It is catalyzed by the protein families of adenosine deaminases acting on RNA (ADAR), which convert adenosine (A) to inosine (I), leading to differences between the RNA products and the corresponding genomic sequences. A-to-I editing is also known as A-to-G editing because inosine is subsequently recognized as guanosine (G) by the cellular translation machinery. Thus, it was suggested that RNA editing could contribute considerably to diversity and flexibility at both the RNA and protein levels, without the need for hardwired mutations at the DNA level (Walkley and Li 2017; Yablonovitch et al. 2017). Although the majority of human RNA editing sites apparently occur in *Alu* repeat elements located in non-coding regions (Athanasiadis et al. 2004; Peng et al. 2012), the number of detected coding editing sites (including synonymous and nonsynonymous sites) is continuing to increase (Tan et al. 2017; Hung et al. 2018). Since nonsynonymous A-to-G RNA editing events can result in amino acid changes at the RNA level (even though most observed coding editing sites are edited at a very low level (Li et al. 2009; Tan et al. 2017)), they may compensate for G-to-A genomic mutations to a certain extent and thus partially reduce the deleteriousness of such mutations. Hence, we are curious about whether nonsynonymous A-to-G RNA editing is associated with the burden of deleterious A/G genomic variants in a population.

To address the abovementioned issue, we conducted the first population-based analysis to examine the association between A-to-G RNA editing activities and the allele frequency of A/G genomic variants in a human population. Since minor alleles tend to be risk alleles (Park et al. 2011; Kido et al. 2018), an advantageous allele should reach a higher allele frequency within a population. If nonsynonymous RNA editing contributes to the prevalence of deleterious A/G genomic changes in a population, there should be a correlation between the editing activity at As and the allele frequency of A. To neutralize the deleterious effect of G-to-A genomic changes at the RNA level, we speculate that As with a lower allele frequency (As tend to be less advantageous than Gs) should be edited more frequently than As with a higher allele frequency (As tend to be more advantageous than Gs). If this is true, then a negative correlation should be observed between A-to-G RNA editing activities (prevalence and level) and the allele frequency of As within a population; meanwhile, this negative correlation should be particularly strong in highly important, evolutionarily conserved loci/genes. However, identification of RNA editing is often hampered by difficulties in distinguishing between true editing sites and A/G genomic variants because of the lack of genome and transcriptome sequencing data from the same samples at the population scale. To minimize false positives arising from single nucleotide polymorphisms (SNPs), sites overlapping with known SNPs (e.g., dbSNP) are generally discarded for detection of RNA editing events (Kurmangaliyev et al. 2015; Walkley and Li 2017; Hung et al. 2018). Thus, such a relationship between A-to-G RNA editing and the allele frequency of A/G genomic variants within a population remains uninvestigated. Recently, the Geuvadis lymphoblastoid cell line (LCL) data (Lappalainen et al. 2013), which encompass both genome and transcriptome sequencing data from 447 individuals, have provided a unique opportunity to determine the RNA editing activities at A/G polymorphic sites within the human population. This dataset provides an ideal resource for our purpose. Moreover, we examined the relationship between the distribution of different SNP types from the human population of apparently healthy individuals (the 1000 Genomes project (Genomes Project et al. 2015)) and the deleterious effects of the corresponding missense changes. We further asked if the relationship is associated with the known sequence preference of ADAR binding (Lehmann and Bass 2000; Eggington et al. 2011) for G depletion and G enrichment at the 5’ and 3’ neighboring nucleotides of the A-to-G editing sites. We thus tested the contribution of nonsynonymous A-to-G RNA editing to the burden of deleterious variants in the healthy human population.

## RESULTS

### Nonsynonymous A-to-G RNA editing activities are associated with the deleteriousness of A/G missense changes

To assess the correlation between coding A-to-G RNA editing activity and the deleteriousness of A/G genomic variants in a population, we first extracted sites with A/G SNPs in coding regions from the Geuvadis LCLs of 447 individuals (derived from the 1000 Genomes project (Genomes Project et al. 2015)) and the RNA-seq data of the corresponding LCL samples from the Geuvadis project (Lappalainen et al. 2013). For each A/G SNP site, there are three possible genotypes across the LCL population: AA, AG, and GG. We selected individuals with the homozygous genotype AA and used the RNA-seq data from the corresponding LCL samples to determine RNA editing (Fig. 1A). Of note, throughout this study we only calculated editing levels at sites with the homozygous genotype AA to eliminate the expression effect of allele G. A site was defined as an editing site if it was found to be edited at a level >5% in at least two LCL samples from individuals with homozygous genotype AA. A total of 1,712 A/G SNP sites satisfied the above rule, in which editing causes a nonsynonymous change at 889 sites and causes a synonymous change at 823 sites (Supplemental File S1).

**Figure 1.**
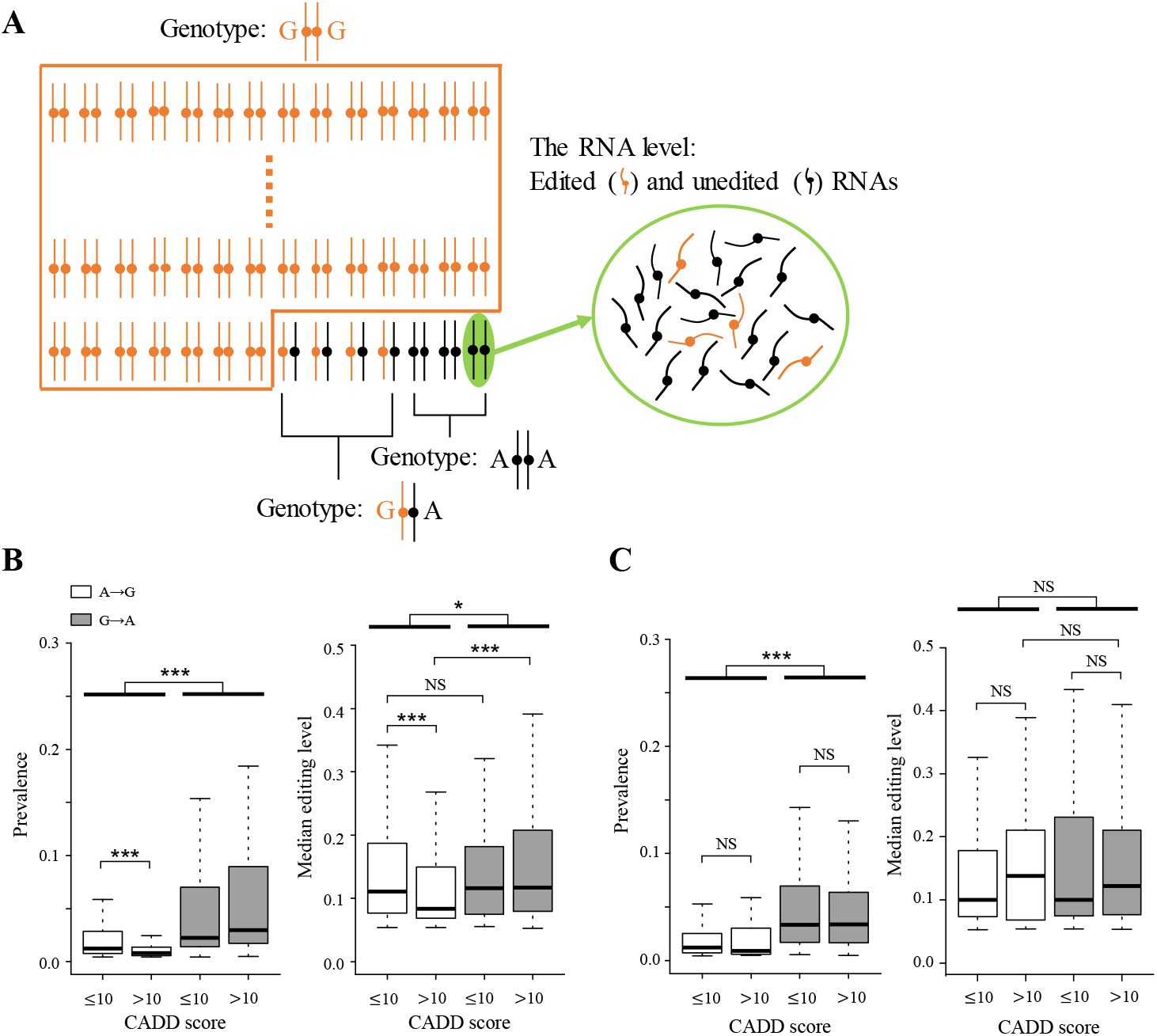
A-to-G RNA editing activities at A/G genomic variant sites. (A) Schematic diagram of an editing event at a variant site with homozygous genotype AA in a population. A and G represent the minor and major alleles in this population, respectively. (B) and (C) Comparisons of (B) nonsynonymous and (C) synonymous editing activities (prevalence and median level) at A/G genomic variant sites with harmless (CADD score ≤ 10) and deleterious (CADD score > 10) A-to-G/G-to-A changes in the LCL population. *P* values were determined using two-tailed Wilcoxon rank-sum test. Significance: \**P* value < 0.05, \*\**P* value < 0.01, and \*\*\**P* value < 0.001. NS, not significant.

We proceeded to examine the correlation between RNA editing activity and the deleterious effect of genomic changes in the population. We used CADD (Kircher et al. 2014), a well-developed tool for measuring the molecular functionality and pathogenicity of genomic changes, to assess the deleteriousness of A/G genomic changes. Intriguingly, two phenomena were observed for the 889 nonsynonymous editing sites (Fig. 1B). First, both the prevalence and level of editing were higher for G-to-A genomic changes than for A-to-G ones. Second, both the prevalence and level of editing were reduced for deleterious (CADD score > 10) A-to-G genomic changes and elevated for deleterious G-to-A changes. Of note, the prevalence of editing at each site was defined as the percentage of individuals with editing done at the site over all individuals with homozygous genotype AA and a read coverage ≥ 10 (i.e., all individuals that were testable for this editing site). Particularly, the median levels of nonsynonymous editing at sites with neutral/harmless (CADD score ≤ 10) A-to-G and G-to-A genomic changes were not statistically different (*P* value = 0.97), whereas the median editing level was significantly higher at sites with G-to-A genomic changes than at those with A-to-G changes when the changes were deleterious (*P* value < 0.001; Fig. 1B, right). In contrast, these phenomena were not observed for synonymous editing sites, although the phenomenon of a higher prevalence of editing for G-to-A genomic changes than for A-to-G ones held (Fig. 1C). These results thus suggest that nonsynonymous RNA editing activities at As are associated with the deleterious effects of A/G missense changes in the human population.

### Nonsynonymous A-to-G RNA editing activities are negatively correlated with the allele frequency of A in a population

We then examined the correlation between nonsynonymous A-to-G RNA editing activity and the allele frequency of A in the LCL population. We first showed that the proportion of deleterious nonsynonymous genomic changes (CADD score > 10) markedly decreased with an increasing minor allele frequency (Supplemental Fig. S1), reflecting the observation that minor alleles tend to be risk alleles (Park et al. 2011; Kido et al. 2018). Next, the above results (Fig. 1B) showed that both the prevalence and level of nonsynonymous editing were negatively correlated with the deleteriousness of A-to-G genomic changes and positively correlated with the deleteriousness of G-to-A genomic changes within a population. Accordingly, for an A/G genomic variant, if G is a minor allele within a population (which implies that G is riskier or less advantageous than A), then the A allele should be edited less to prevent the conversion of A into I (which is then recognized as G) at the RNA level. In contrast, if A is a minor allele (which implies that A is riskier or less advantageous than G), then editing of the A allele should be promoted to compensate to a certain extent for the deleterious G-to-A change. In other words, we speculated that both the prevalence and level of nonsynonymous editing should negatively correlate with the allele frequency of A within the LCL population. Indeed, we observed that both the prevalence (Fig. 2A) and level (Fig. 2B) of nonsynonymous RNA editing markedly decreased with an increasing allele frequency of A (or with a decreasing allele frequency of G) within the LCL population. Again, this result supports the correlation between nonsynonymous RNA editing activities and the deleteriousness of A/G missense changes in the human population.

**Figure 2.**
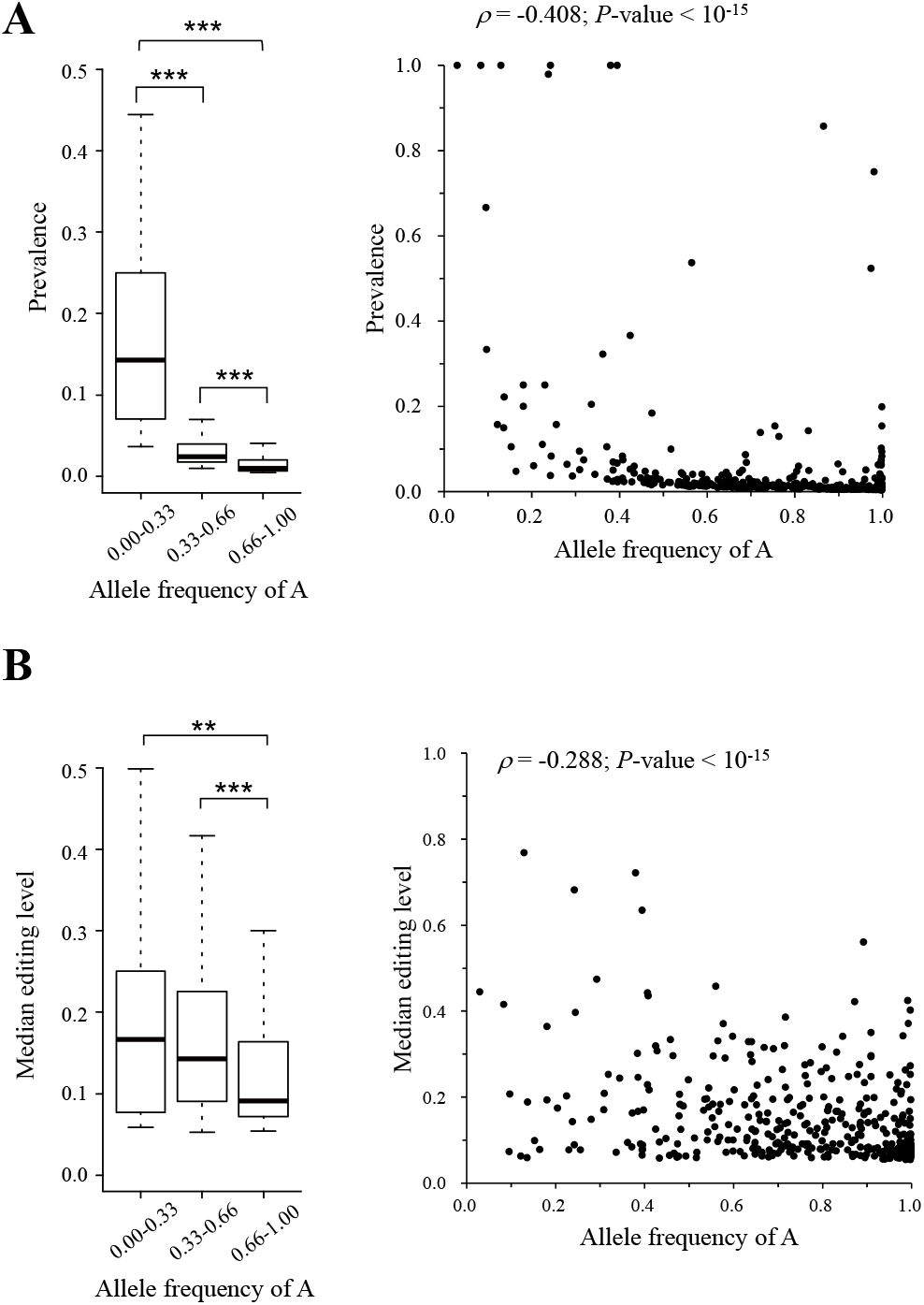
Correlation between nonsynonymous editing activities and allele frequency of A. Box (left) and scatter (right) plots represented the correlations between nonsynonymous editing activities ((A) prevalence and (B) median editing level) and allele frequency of A within the LCL population. Significant differences between prevalence/median editing level and allele frequency of A were determined using two-tailed Wilcoxon rank-sum test. Significance: \*\**P* value < 0.01 and \*\*\**P* value < 0.001.

### The negative RNA editing-allele frequency correlation is stronger in functionally more important loci/genes than in less important ones

We have observed a significant correlation between nonsynonymous RNA editing activities and the deleteriousness of A/G missense changes in a population. We then asked whether such a correlation was stronger in functionally more important loci/genes than in less important ones. If it was, we should observe a stronger correlation between nonsynonymous RNA editing activities and allele frequency of A within a population in functionally more important loci/genes than in less important ones. To this end, we performed the following evolutionary and functional analyses (see Figs. 3A and 3B and Supplemental Fig. S2). First, we considered the selective constraints of the host genes that contained the nonsynonymous editing sites examined. We divided nonsynonymous editing sites into two equally sized groups (i.e., editing sites located within genes under strong and weak selection pressure) according to the ratios of nonsynonymous to synonymous substitution rates (*dN*/*dS*) calculated using one-to-one human-rhesus macaque and human-mouse orthologs, respectively. Indeed, we found that the negative correlations between nonsynonymous editing activities and the allele frequency of A were significantly stronger for editing sites located within genes under strong selective constraints than for those located within genes under weak selective constraints (all *P* values < 0.001 by the two-tailed Z score test). Second, we examined whether this correlation was affected by the conservation of individual nucleotides (as determined by PhyloP (Pertea et al. 2011) and PhastCons (Siepel et al. 2005) scores). Similar to the above analysis, we divided the nonsynonymous editing sites into two equally sized groups: highly and lowly conserved sites. As expected, the correlations were significantly stronger for highly conserved sites than for lowly conserved ones. Third, we examined the effect of gene essentiality on the correlation because essential genes are those that are functionally indispensable for the survival of an organism. We divided nonsynonymous editing sites into two groups: sites located within human essential (including essential and conditional essential) genes and those located within non-essential genes. Of note, gene essentiality information was retrieved from the OGEE database, which comprises 16 gene essentiality data sets derived from nine human cancers (Chen et al. 2017). We also found a significantly stronger negative correlation between the prevalence/level of nonsynonymous editing and the allele frequency of A for editing sites located within essential genes than for those located within non-essential ones. Fourth, we further predicted that such a negative correlation should be stronger for editing sites located within genes intolerant of a loss of function (LOF) mutation than for those located within genes with LOF tolerance. We considered gene variant intolerance (pLI) scores estimated by the ExAC Consortium (Lek et al. 2016), which range from 0 to 1 with higher pLI scores indicating greater LOF intolerance levels. Human genes were divided into two groups using the 90th percentile as a cut-off. The resulting data, shown in Figures 3A and 3B, strongly supported our prediction. Finally, we examined the effect of gene pleiotropy on the correlation. Pleiotropy is a phenomenon in which a single gene can result in multiple distinct phenotypes. Pleiotropic genes were reported to be essential, to be highly connected in protein-protein interaction networks, and to highly contribute to human diseases (Ittisoponpisan et al. 2017). Generally, we found more strongly negative correlations between nonsynonymous editing activities (editing level, particularly) and the allele frequency of A for editing sites located within pleiotropic genes than for those located within non-pleiotropic ones. Taken together, these results support that the negative RNA editing-allele frequency correlation is stronger in functionally more important loci/genes than in less important ones in the human population.

**Figure 3.**
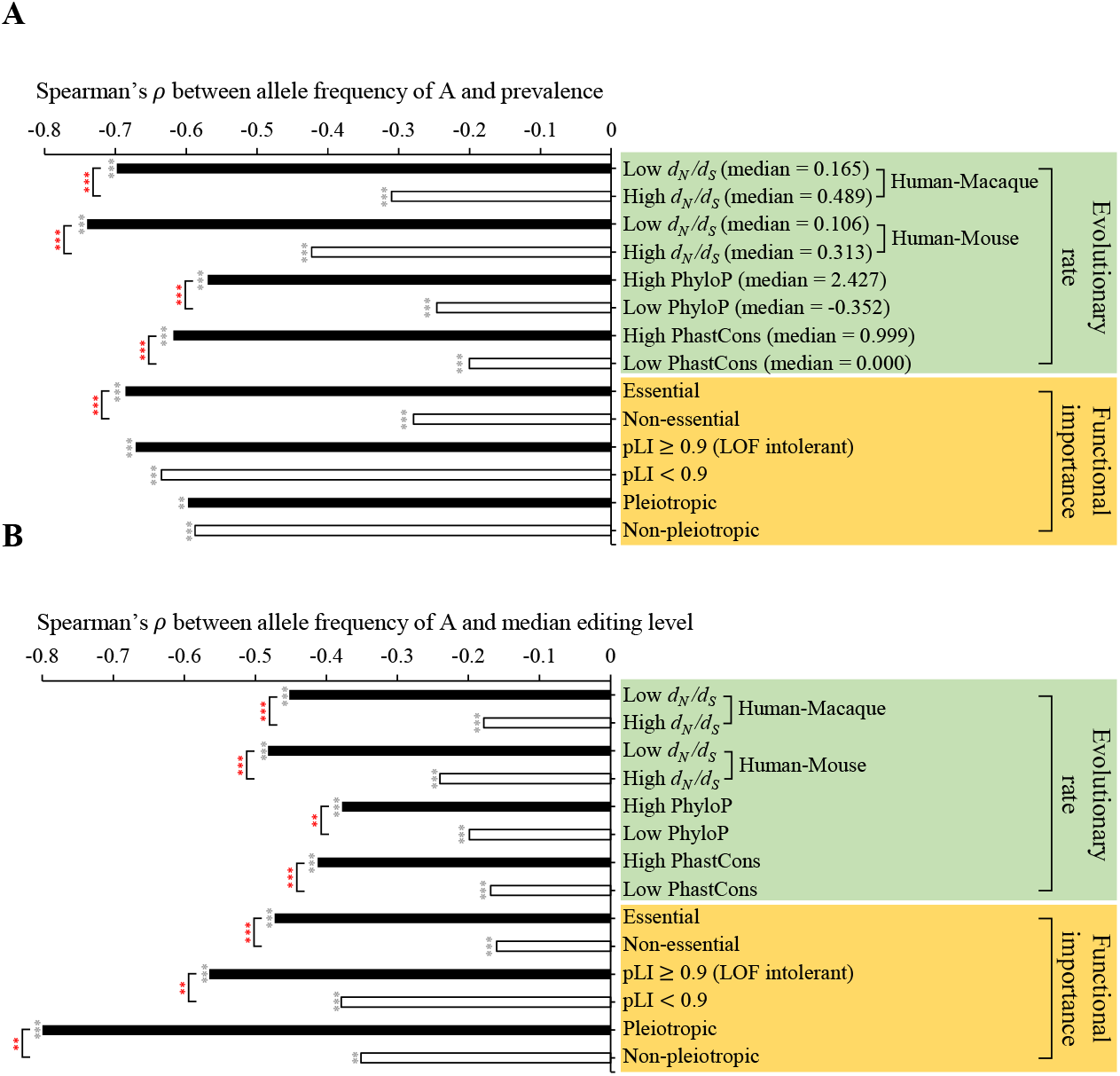
Functional and evolutionary analysis of the RNA editing-allele frequency correlations. The histograms represented the correlations between nonsynonymous editing activities ((A) prevalence and (B) median editing level) and allele A frequency within the LCL population in four categories of evolutionary rates and three categories of functional importance (see the text) of the target genes/loci where the editing sites were located. For four categories of evolutionary rates, editing sites were divided into two equal groups according to the high and low scores of the target genes/loci. The statistical significances of Spearman’s rank correlation coefficients (*ρ*) were represented by black stars. Significant differences between two independent correlations (represented by red stars) were estimated using two-tailed Z score test with the paired.r function within the *psych* R library. Significance: \**P* value < 0.05, \*\**P* value < 0.01, and \*\*\**P* value < 0.001.

### Phylogenetic variation in the RNA editing-allele frequency correlation

The pattern of phylogenetic variation of a trait is often used to assess its functional importance. Regarding the patterns of phylogenetic variations for the 889 nonsynonymous editing sites, we can assess the RNA editing-allele frequency correlation in different phylogenetic types of nonsynonymous A-to-G editing sites. To this end, we retrieved human, chimpanzee, rhesus macaque, and mouse orthologous nucleotides and defined five types of human nonsynonymous editing sites according to their phylogenetic variations (Fig. 4A). First, “G-conserved” sites were those for which a G allele was observed in all three non-human orthologs. Second, “A-conserved” sites were those for which an A allele was observed in all three non-human orthologs. Third, “hardwired” sites were those for which either an A or G was observed in all three non-human orthologs. Fourth, “G-unfound” sites were those for which G was not observed in any of the three non-human orthologs, excluding sites already designated as A-conserved. Fifth, “diversified” sites were those for which either a G or non-G was observed in all non-human orthologs, excluding sites already designated as hardwired. This categorization resulted in 100 G-conserved sites, 299 A-conserved sites, 114 hardwired sites, 35 G-unfound sites, and 26 diversified sites. Of note, A/G coincident SNPs (Hodgkinson et al. 2009; Chen et al. 2016), which were orthologous sites observed to be A/G polymorphic in any two examined species, were considered as the hardwired sites. Regarding the patterns of phylogenetic variations, diversified sites may be subject to the weakest selective constraints compared to other types of sites because these positions allow various amino acid changes (Xu and Zhang 2014). Thus, we predicted that the RNA editing-allele frequency correlation should be the weakest at diversified sites because editing would cause the smallest impact on these sites, given their tolerance for variation in amino acids. Indeed, we found that the correlation between the median editing level and the allele frequency of A at diversified sites was insignificant (*P* value = 0.12); in contrast, the negative RNA editing-allele frequency correlations held significantly for all the other types of sites (Fig. 4B and Supplemental Fig. S3).

**Figure 4.**
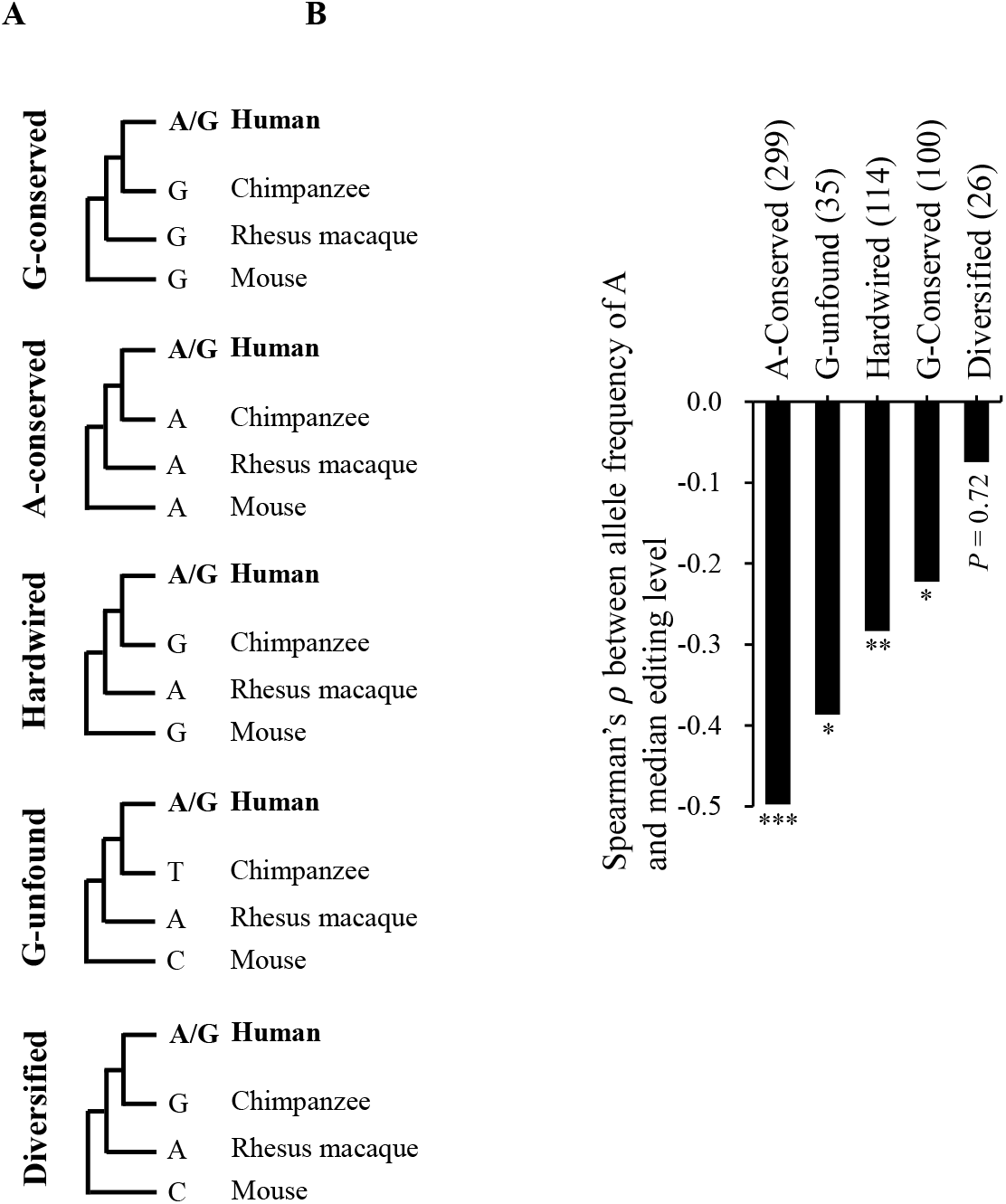
Phylogenetic analysis of nonsynonymous A-to-G editing sites. (A) Definition of five types of human nonsynonymous editing sites (A-conserved, G-unfound, hardwired, G-conserved, and diversified) based on their evolutionary variations among human, chimpanzee, rhesus macaque, and mouse. (B) Correlations between median editing levels and allele frequency of A within the LCL population for the five groups of editing sites. The number of nonsynonymous editing sites examined in each group was provided in parentheses. *P* values were determined using two-tailed Wilcoxon rank-sum test. Significance: \*\**P* value < 0.01 and \*\*\**P* value < 0.001.

### Deleterious G-to-A missense changes are more tolerable than other types of changes in a population

The above results that nonsynonymous editing activities are correlated with the deleteriousness of A/G genomic changes raise the interesting question of whether nonsynonymous editing is associated with the distribution of missense variants (or nonsynonymous SNPs) in a population, especially when the missense changes are damaging. To this end, we extracted rare missense mutations (Materials and Methods) from the 1000 Genomes project (Genomes Project et al. 2015) and examined the correlation between the distribution of six possible SNP types and the deleterious effects of the corresponding missense changes (Supplemental Table S1). The SNPs were classified into six groups rather than twelve because genomic variants could potentially cause variants in both the sense and antisense transcripts (e.g., A-to-G changes on one strand and T-to-C changes on the opposite strand). It was not surprising that there was a higher proportion of rare missense mutations in the two transition groups (A-to-G/T-to-C and G-to-A/C-to-T changes) than in the four transversion groups (Fig. 5A). Interestingly, the two transition groups exhibited quite different trends in terms of the proportions of rare missense mutations. Proportions of rare missense mutations were markedly negatively correlated with the corresponding CADD scores for A-to-G/T-to-C changes, but the reversed trend was observed for G-to-A/C-to-T changes (Fig. 5A). In particular, regarding the very deleterious missense changes (e.g., CADD scores > 30), the vast majority (84%) of rare missense variants were G-to-A/C-to-T changes while only 2% were A-to-G/T-to-C changes (Fig. 5A). The result suggests that G-to-A/C-to-T missense changes are more acceptable than other types of changes in the human population when the changes are deleterious. Considering all human coding regions, we further compared the frequencies of two types of transitions (i.e., *f*(G-to-A/C-to-T) vs. *f*(A-to-G/T-to-C)) in different CADD scores (see Materials and Methods and Supplemental Table S1). We found that the *f*(G-to-A/C-to-T)/*f*(A-to-G/T-to-C) ratio was relatively close to one when the rare missense changes were neutral/harmless (CADD score ≤ 10); however, such ratios were significantly higher than one when the changes were deleterious (CADD score > 10) (Fig. 5B). Particularly, the *f*(G-to-A/C-to-T)/*f*(A-to-G/T-to-C) ratios markedly increased with increasing CADD scores (Fig. 5B). Moreover, we calculated the proportions of rare missense mutations for A-to-G/T-to-C and G-to-A/C-to-T changes per individual from the 1000 genome project and examined the individual mutational burden of these two types of rare missense changes. We also found that the median proportion of G-to-A/C-to-T missense changes was significantly higher than that of A-to-G/T-to-C changes, particularly when the changes were deleterious (Fig. 5C, upper). The differences in mutational burden between these two types of rare missense changes markedly increased with increasing deleterious effects of the changes (Fig. 5C, bottom). These results further support that damaging G-to-A/C-to-T missense mutations are especially tolerable in a healthy population.

**Figure 5.**
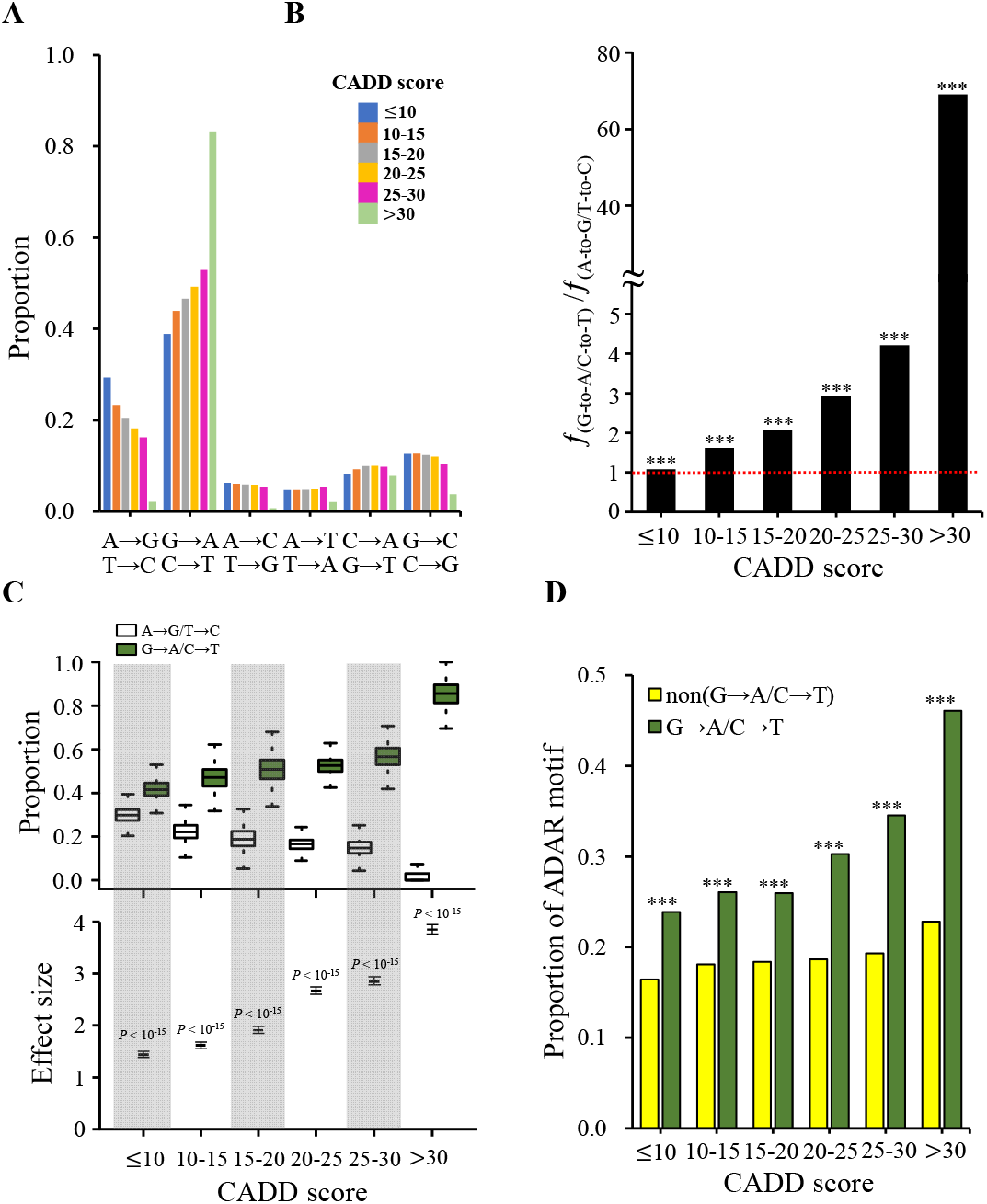
Relationship between the distributions of different types of rare missense variants and A-to-G RNA editing. (A) Correlation between the distributions of different types of rare missense variants from the 1000 genome project and deleteriousness (measured by CADD scores) of the corresponding genomic changes. (B) Fraction of the frequency of nonsynonymous G-to-A/C-to-T changes to the frequency of nonsynonymous A-to-G/T-to-C changes (i.e., the *f*_(G-to-A/C-to-_ T)/*f*_(A-to-G/T-to-C)_ ratio) for different deleterious effects of the genomic changes. (C) Comparisons of the individual mutational burden of the two types of rare transition missense variants (A-to-G/T-to-C and G-to-A/C-to-T) in the 1000 genome project. The upper panel represented the proportions of rare A-to-G/T-to-C and G-to-A/C-to-T missense variants per individual for different CADD scores. The bottom panel represented the differences (effect sizes) mutational burden between these two types of rare missense variants. Effect sizes were measured by Cohen’s *d*, which was defined as the difference between both mean numbers of these two types of rare missense variants divided by the standard deviation of the paired differences. The estimated 95% confidence intervals of effect sizes were plotted (see also Supplemental Table S2). (D) Comparisons of the proportions of SNP sites with the ADAR motif for different deleterious effects of G-to-A/C-to-T and non-G-to-A/C-to-T rare missense variants. *P* values were determined using two-tailed Fisher’s exact test ((B) and (D)) or two-tailed Wilcoxon signed ranked test (C). Significance: \*\**P* value < 0.01 and \*\*\**P* value < 0.001.

We then examined whether the preference for damaging G-to-A/C-to-T missense mutations in a population was associated with the primary ADAR sequence motif (Lehmann and Bass 2000; Eggington et al. 2011) for G depletion and G enrichment at the 5’ and 3’ neighboring nucleotides of the A-to-G editing sites. Indeed, sites of G-to-A/C-to-T missense mutations have a significantly higher percentage of the ADAR motif than those of non-G-to-A/C-to-T ones; such differences markedly increased with increasing CADD scores (Fig. 5D). All the trends observed above are independent of tools for measuring the deleteriousness of amino acid substitutions (Supplemental Fig. S4) and human subpopulations (Supplemental Fig. S5). These observations thus suggest that nonsynonymous A-to-G RNA editing is highly associated with the distribution of existing nonsynonymous polymorphisms at functionally important loci, contributing to the mutational burden of deleterious missense variants in the human population.

## DISCUSSION

This study utilized the Geuvadis LCL data, which provide both genome and transcriptome sequencing data from the same individuals at a population scale, and conducted the first population-based analysis to examine the relationship between A-to-G RNA editing activities and the allele frequency of A/G genomic variants. Our results revealed that, for existing A/G missense variants, both the prevalence and level of nonsynonymous A-to-G RNA editing were higher at sites with deleterious G-to-A genomic changes than at those with deleterious A-to-G ones (Fig. 1B) and that nonsynonymous editing activities exhibited a significantly negative correlation with the allele frequency of A in the LCL population (Fig. 2). The negative editing-allele frequency correlation is significantly stronger in functionally more important loci/genes than in less important ones (Fig. 3); of note, such a trend is observed for nonsynonymous editing, but not for synonymous editing when considering the correlations between editing level and allele frequency of A (Supplemental Fig. S6). We further observed that G-to-A/C-to-T missense mutations were much more prevalent than A-to-G/T-to-C changes (and other types of changes) at functionally important sites, in which the differences in the mutational burden of missense changes was significantly positively correlated with the deleteriousness of the changes (see Figs. 5A, 5B, and 5C). For an existing A/G missense variant, if the G-to-A genomic change has severely deleterious effects on protein function, RNA editing at this site with a higher editing level is more likely to neutralize the deleterious effect of the genomic change at the RNA level, making the deleterious effect weaker than expected. RNA editing may facilitate the escape of this existing deleterious A/G variant from negative natural selection and therefore aid the tolerance for this deleterious variant (i.e., the deleterious allele A) in a population. We thus suggest that nonsynonymous A-to-G RNA editing is associated with the missense variant distribution and contributes to mutation load in a population.

Since it is known that DNA methylation at CpG dinucleotides can significantly increase the rate of spontaneous C-to-T transitions (Coulondre et al. 1978; Bird 1980; Holliday and Grigg 1993), one may ask whether DNA methylation might affect the preference for G-to-A/C-to-T missense mutations at functionally important loci in a population. In human, it was observed that the majority of gene bodies were heavily methylated (Keller et al. 2016). Position-dependent correlations have also been shown between CpG methylation level and nonsynonymous substitution rate (*dN*) of the target genes (Chuang and Chiang 2014) or exons (Chuang et al. 2012); such methylation-*dN* correlations were observed to be negative for gene bodies (or internal/last exons) and positive for promoter regions (or first exons). Accordingly, we asked whether our results shown in Figure 5 were biased toward first exons. We divided nonsynonymous SNP sites into two groups: sites located within the first and non-first (internal/last) exons. We performed the similar analysis and found that all the trends observed in Figure 5 held well in both first and non-first exons (Supplemental Fig. S7), suggesting that the effect of CpG methylation is less likely to be responsible for the preference for damaging G-to-A/C-to-T missense mutations in a population. Meanwhile, although methylation level is positively correlated with *dN* of first exons (Chuang et al. 2012), methylation effect seems not to be the cause of this preference in first exons for the following two reasons. First, the proportion of lowly methylated promoters is markedly higher than that of heavily methylated ones in human (Keller et al. 2016), although the bimodal pattern of lowly and heavily methylated promoters is generally observed in vertebrates (Saxonov et al. 2006; Elango and Yi 2008; Lou et al. 2014; Mendizabal and Yi 2016). Second, for nondegenerate nucleotides (i.e., all genomic changes at these sites must cause nonsynonymous changes) in first exons, methylated nucleotides are more conserved than unmethylated ones when their target exons are subject to stringent selective constraints (Chuang and Chen 2014). These observations thus suggest that the effect of DNA methylation on coding regions is unlikely to be a consequence of the preference for damaging G-to-A/C-to-T missense variants in the human population.

That nonsynonymous editing may help tolerate existing deleterious A/G variants in a population raises the question of whether RNA editing is generally adaptive. Several studies have speculated that nonsynonymous editing is more beneficial for enhancing transcriptome (and therefore proteome) diversity and fitness than the direct replacement of As with Gs at the DNA level, and thus it is maintained by natural selection (Gommans et al. 2009; Li et al. 2009; Nishikura 2010; Porath et al. 2017b; Eisenberg and Levanon 2018). However, most A-to-G RNA editing events are considered to originate through promiscuous targeting by ADAR proteins (Xu and Zhang 2014). Like missense mutations, most nonsynonymous A-to-G RNA editing events may result in deleterious effects on protein function and consequently be eliminated from a population by natural selection. Several observations support the claim that editing events are generally damaging. First, human coding RNA editing events are rare and tend to be synonymous (Kleinman et al. 2012; Chen 2013). Second, only a small number of conserved mammalian editing sites have been observed (Pinto et al. 2014; Liscovitch-Brauer et al. 2017). Third, editing activity generally exhibits a minor quantitative impact on highly expressed genes, essential genes, or regions under strong selective constraints (Solomon et al. 2014; Xu and Zhang 2014). Finally, edited As were more likely to be replaced with Gs than unedited As in evolution (Xu and Zhang 2014). Taken together, although natural selection has distinctly shaped the landscapes of the editomes in various phylogenetic clades (Yablonovitch et al. 2017), human RNA editing does not exhibit the signals of adaptation overall.

In addition, according to phylogenetic analysis of some specific cases, RNA editing was suggested to be advantageous by extending sequence divergence at the DNA level, acting as a safeguard by correcting G-to-A mutations at the RNA level and thus mediate the RNA memory of evolution (Tian et al. 2008; Chen 2013). However, the safeguard scenario is unlikely to occur in functionally important regions of the genome because most nonsynonymous A-to-G RNA editing events are edited at a very low level (Solomon et al. 2014). It was observed that the level of coding editing was generally less than 10% (Tan et al. 2017) except for very few events of high editing levels (e.g., the editing site at position 607 of *GRIA2* (Sommer et al. 1991); or see the summary table in Hung el al.’s study (Hung et al. 2018)). Most nonsynonymous editing events cannot fully post-transcriptionally compensate for the deleterious effects of G-to-A genomic mutations. In addition, regarding the five phylogenetic types of nonsynonymous A-to-G editing (Fig. 4A), we observed that the median editing level of diversified sites is similar to that of G-conserved and hardwired sites (both *P* values > 0.05 by two-tailed Wilcoxon rank-sum test; Supplemental Fig. S8A). Diversified sites are tolerant of many different amino acids; in contrast, Gs at both G-conserved and hardwired sites are under relatively stronger selective constraints. If A-to-G editing is mostly advantageous and acts as a safeguard of G-to-A substitutions, then the editing level should be significantly higher at G-conserved and hardwired sites than at diversified sites. Again, this observation supports the above discussion that nonsynonymous editing is generally nonadaptive and unlikely to have a safeguard role overall in human.

Moreover, we found that the median editing level at A-conserved and G-unfound sites was significantly lower than the levels at diversified sites (Supplemental Fig. S8A). Since Gs at both A-conserved and G-unfound sites are not selectively permitted during mammalian protein evolution, most editing events at genomic As should be constrained. This result is consistent with a previous study (Xu and Zhang 2014), which performed a similar analysis by categorizing human nonsynonymous editing sites into four phylogenetic types (i.e., A-conserved, hardwired, G-unfound, and diversified sites) based on phylogenetic variations among 44 non-human vertebrates (Supplemental Fig. S8B). This observation also reflects our abovementioned results that nonsynonymous A-to-G RNA editing activities are significantly reduced at A/G missense variants when A-to-G genomic changes are deleterious (Fig. 1B).

In conclusion, this study highlights the association between nonsynonymous RNA editing and existing missense variants in the human population. Although the cause and distribution of damaging variants within a population are more complicated than expected (Henn et al. 2015), our findings reveal that nonsynonymous A-to-G RNA editing is associated with the increased burden of deleterious G-to-A missense variants in the healthy human population. Particularly for pathogenomics studies, deleterious variants are often observed in well-established disease-associated genes in population controls. Our results thus call to pay special attention to nonsynonymous A-to-G RNA editing in pathogenomics studies for extracting pathogenic variants.

## METHODS

### Identification of RNA editing sites in the LCL population

The human gene annotation and the strands of sites were downloaded from the Ensembl genome browser at http://www.ensembl.org/ (version 87). The genotype data were extracted from the Geuvadis LCLs of 447 individuals [derived from the 1000 Genomes project (Genomes Project et al. 2015)]. The RNA-seq data of the corresponding LCL samples were extracted from the Geuvadis project at https://www.ebi.ac.uk/Tools/geuvadis-das/ (Lappalainen et al. 2013). The maskOutFa tool (https://github.com/ENCODE-DCC/kentUtils/tree/master/src/hg/maskOutFa) was used to mask the SNP sites of corresponding LCL samples at the human reference genome (GRCh38.p7) and generate the pseudo-genome for each individual. The corresponding RNA-seq reads were then aligned against the pseudo-genome using STAR aligner (version 2.5.1b) (Dobin et al. 2013). For the purpose of studying the relationship between allele frequency of A/G genomic variants and A-to-G RNA editing, we only considered sites with A/G polymorphisms across the LCL population. There are three possible genotypes for an A/G SNP site: AA, AG, and GG. We used SAMtools (version 1.2) mpileup (Li 2011) and a pileup parser program pileup_parser.pl to call variants at the A/G SNP sites with the homozygous genotype AA for each individual. Of note, to eliminate the expression effect of allele G, throughout this study we only calculated editing levels at sites with the homozygous genotype AA. The editing level of a site was determined by the ratio of number of G reads to the sum of numbers of A and G reads (i.e., RNA editing level = G / (A+G)). We only simultaneously considered the sites with ≥ 10 matched reads. A site was defined as an editing site if it satisfied two rules: (1) the base quality score must be ≥ 25 with the STAR-mapping quality score (Dobin et al. 2013) ≥ 255 and (2) the site was found to be edited at a level >5 % in at least two LCL samples from individuals with homozygous genotype AA. To minimize potential false positives caused by sequencing errors, an editing event was not considered if its editing level was ≥ 90%. The identified editing events and the related data were deposited in Supplemental File S1.

### Deleterious effects of genomic changes

The deleteriousness of genomic changes was evaluated by Combined Annotation Dependent Depletion (CADD) score (Kircher et al. 2014), which can quantitatively prioritize the functional categories for wide range variants, such as functional, deleterious, and disease causal variants. The CADD score is a Phred-scale value, with higher CADD scores indicating that the genomic changes had more damaging effects. For the genomic changes from the 1000 Genome project, the CADD scores of the changes were downloaded directly from the 1000 Genome project. For all possible nonsynonymous genomic changes in all human coding regions, the CADD scores of the changes were downloaded from the CADD browser (release v1.4) at http://cadd.gs.washington.edu/. From the 1000 genome project, we extracted rare missense mutations from derived alleles (based on the chimpanzee reference genome) with minor allele frequency less than 0.01. We took G-to-A/C-to-T rare missense mutations as an example to describe the calculation of frequencies of the genomic changes with different deleterious effects in the 1000 Genome project. As shown in Supplemental Table S1, there are 40,239 G-to-A/C-to-T rare mutations that cause nonsynonymous changes with a CADD score ≤ 10 in the 1000 Genome project. Of all human coding regions, 690,353 G (or C) sites would have nonsynonymous changes with a CADD score ≤ 10 if changed to A (or T). The frequency of G-to-A/C-to-T rare missense mutations with a CADD score ≤ 10 is *f*(G-to-A/C-to-T) = 40,239/690,353 = 5.83×10^-2^.

### Evolutionary and functional analysis

The *dN/dS* ratios between human-rhesus macaque orthologs and between human and mouse orthologs were downloaded from the Ensembl genome browser (version 87). The lower *dN/dS* ratios indicate that the examined genes have stronger selective constraints. Both PhyloP (Pertea et al. 2011) and PhastCons (Siepel et al. 2005) scores were downloaded from the UCSC genome browser at http://ge-nomes.ucsc.edu/. The higher PhyloP/PhastCons scores indicate higher conservation levels of the examined sites. The essentiality of human genes was obtained from the Online GEne Essentiality (OGEE) (Chen et al. 2017) database. For human, the OGEE database consists of 183 essential, 6,985 conditionally essential, and 14,388 non-essential genes. OGEE essential and conditionally essential genes were both considered as essential genes for this study. The gene variant intolerance (pLI) scores were estimated by the Exome Aggregation Consortium (ExAC), which range from 0 to 1, with higher pLI scores indicating greater LOF intolerance levels (Lek et al. 2016). The pLI scores of human genes were retrieved from the ExAC browser at http://exac.broadinstitute.org/. Human genes were divided into two groups, 3,488 LOF-intolerant and 14,753 LOF-tolerant genes, using the 90th percentile as a cutoff. The information on 412 pleiotropic and 1,345 non-pleiotropic genes was downloaded from PleiotropyDB (Ittisoponpisan et al. 2017) at http://www.sbg.bio.ic.ac.uk/pleiotropydb/home/. Genes were classified as pleiotropic when they were associated with more than one disorder affecting different physiological systems or different conditions of a single physiological system, whereas genes were classified as non-pleiotropic when they were associated with only one disorder (Ittisoponpisan et al. 2017).

### Phylogenetic variation of nonsynonymous editing sites

We retrieved the pattern of phylogenetic variation for each analyzed nonsynonymous editing site from the human, chimpanzee, rhesus macaque, and mouse orthologous nucleotide. The corresponding genome coordinates of orthologous sites among these species were determined using the UCSC liftOver tool (https://genome.ucsc.edu/cgi-bin/hgLiftOver). An orthologous site was determined as an A/G co-incident SNP if it was observed to be A/G polymorphic in any two examined species. The SNP information on these non-human species were downloaded from the Ensembl database (version 87).

### Statistical analysis

Statistical analyses were performed using R software version 3.4.2 (http://www.r-project.org/). Statistically significant differences were determined using two-tailed Wilcoxon rank-sum test, two-tailed Wilcoxon signed ranked test, two-tailed Fisher’s exact test, and Spearman’s rank correlation, as appropriate. Statistically significant difference between two independent correlations were estimated from two-tailed Z score test using the paired.r function within the *psych* R library.

### Data access

Datasets generated and analyzed during the current study are presented in the paper and/or the supplementary materials (Supplemental File S1).

## Acknowledgments

We thank Chia-Ying Chen for discussions and Yu-Shiang Zeng for computational assistance. This work was supported by Genomics Research Center (GRC), Academia Sinica, Taiwan; and the Ministry of Science and Technology, Taiwan, under the contract MOST 103-2628-B-001-001-MY4 (to T.-J. C.).

